# Dopaminergic axon tracts within a hyaluronic acid hydrogel encasement for implantation to restore the nigrostriatal pathway

**DOI:** 10.1101/2021.07.03.451006

**Authors:** Wisberty J. Gordián-Vélez, Kevin D. Browne, Jonathan H. Galarraga, John E. Duda, Rodrigo A. España, H. Isaac Chen, Jason A. Burdick, D. Kacy Cullen

## Abstract

Parkinson’s disease (PD) affects 10 million patients worldwide, making it the second most prevalent neurodegenerative disease. Motor symptoms emerge from the loss of dopamine in the striatum after the death of dopaminergic neurons and the long-projecting axons of the nigrostriatal pathway. Current treatments, such as dopamine replacement, deep brain stimulation or cell therapies, disregard the loss of this pathway at the core of symptoms. We sought to address this by improving our tissue-engineered nigrostriatal pathway (TE-NSP) technology, which consists of a tubular hydrogel with a collagen/laminin core that encases an aggregate of dopaminergic neurons and their axons in a way that resembles the nigrostriatal pathway. These constructs can be implanted to replace the lost neurons and axons with fidelity to the pathway, and thus provide dopamine according to feedback from the host circuitry. While TE-NSPs have been traditionally fabricated with agarose, here we utilized a hyaluronic acid (HA) hydrogel to expand the functionality of the encasement and our control over its properties. Using rat ventral midbrain neurons, we found that TE-NSPs exhibited longer and faster neurite growth with HA relative to agarose, with no differences observed in electrically-evoked dopamine release. When transplanted, HA hydrogels reduced host neuron loss and inflammation around the implant compared to agarose, and the cells and axons within TE-NSPs survived and maintained their cytoarchitecture for at least 2 weeks.

**Highlights:**

- We fabricated engineered dopaminergic axons encased in a tubular hydrogel.
- We made hydrogels from methacrylated hyaluronic acid and compared them to agarose.
- Axons in HA hydrogels had longer and faster axon growth and displayed evoked dopamine release.
- HA hydrogels reduced the host inflammatory response and supported neuron and axon survival *in vivo*.
- This platform may be used to reconstruct the nigrostriatal pathway to treat Parkinson’s disease.

## Introduction

Parkinson’s disease (PD) is the second most common neurodegenerative disease in the world, and its main symptoms involve motor deficits such as tremors, bradykinesia, and rigidity [1,2]. These emerge from the degeneration of the nigrostriatal pathway, which consists of dopaminergic neurons, located in the substantia nigra pars compacta, that project long axons that synapse with the striatum and release the dopamine necessary to properly regulate motor function. A standard treatment for PD is dopamine replacement with drug therapies (*e.g.*, L-DOPA), which generally involve discontinuous drug delivery that can cause severe motor fluctuations with chronic exposure [2]. Another common strategy is deep brain stimulation (DBS), where stiff inorganic electrodes are implanted downstream of the striatum; however, DBS does not actually address the causes of PD [2]. As such, several preclinical and clinical studies have looked at the transplantation of fetal- or stem cell-derived dopaminergic neurons in the striatum to restore innervation and the lost dopamine input [3]. Although cell therapies move away from purely symptom management and attempt to address some of the underlying causes of PD, such approaches still neglect reconstructing the nigrostriatal pathway and disregard the correct anatomical location of these neurons and their long projections.

The anatomy of the nigrostriatal pathway has functional implications, as it enables the regulation of dopaminergic neurons by other regions of the basal ganglia and permits synaptic connections between these neurons and adjacent areas [4,5]. This closed loop is crucial to support feedback mechanisms for proper dopamine release in the striatum. Because of this, some studies have involved cellular implants in the substantia nigra in an attempt to promote long-distance axon growth *in vivo* to restore the pathway and reinnervate the striatum with greater anatomical fidelity [6,7]. These studies have resulted in varying degrees of success and are limited by instances of uncontrolled *in vivo* growth and, in some cases, by inherent impediments to clinical translation. Another concern with cellular implants is that these strategies typically employ cells cultured on two-dimensional (2D) surfaces and delivered as suspensions into the striatum, either manually or by infusion pump with a needle injection [8–11]. Regrettably, the vast majority of delivered cells die after implantation, with studies reporting cell losses averaging 92-99% [3,12]. Such severe cell death may be attributed to the stress associated with injections, the transition from 2D culture to the three-dimensional (3D) brain tissue, a lack of extracellular matrix, mass transport limitations, and host inflammatory responses [13]. The field of biomaterials has responded in part with hydrogels for 3D cell encapsulation, given that interactions between cells and their 3D matrix are paramount to activate signaling pathways related to survival, growth, differentiation, and integration [12]. In addition, the properties of these hydrogels may be tuned to provide a substrate for adhesion, promote survival during transplantation, shield cells from inflammation, and provide inductive growth factors [13,14]. For example, encapsulation of neural progenitors and stem cell-derived neurons has led to significant improvements in survival, functional recovery, and reduced graft-induced inflammation in stroke models, and this augmented viability was linked to receptor binding to the hydrogel [12,14]. Recent studies have encapsulated human dopaminergic neurons for both *in vitro* 3D differentiation and implantation, showing a 5-fold enhancement of survival relative to suspensions of cells cultured on 2D surfaces [15,16]. Regardless, in these studies the surviving cells were still a fraction of the implanted ones, particularly at chronic time points. Despite the need for further research to maintain high cell viability, these studies highlight the importance of properly culturing and protecting cells for implantation.

Our group has developed a platform that (1) addresses the limitations of common treatment strategies for PD, (2) reconstructs both the neurons and axon tracts of the nigrostriatal pathway, (3) combines cell replacement with tissue engineering, and (4) employs a tubular hydrogel encasement for 3D axon growth *in vitro* and protection *in vivo*. This platform, called the tissue-engineered nigrostriatal pathway (TE-NSP), consists of aggregates of rat embryonic ventral midbrain neurons that project aligned axon tracts within a hydrogel micro-column containing an extracellular matrix (ECM) core, which enables survival and growth [17]. These constructs are initially fabricated *in vitro* to structurally and functionally mimic the nigrostriatal pathway **(Fig. 1)** and are then implanted to span this pathway and replace both the neurons and axons [17]. TE-NSPs may also be used as *in vitro* test beds to increase our understanding of human midbrain development and the pathophysiology of PD [18]. TE-NSPs fabricated with >50% dopaminergic neurons produced axons growing >5 mm after 21 days *in vitro* (DIV), physically integrated with an aggregate of striatal neurons, released dopamine after electrical stimulation, and survived and grew into the striatum when implanted for 1 month in a rat PD model [19]. These constructs have been made traditionally with agarose, a biomaterial that is not adhesive to mammalian cells and is incapable of significant degradation *in vivo* [20–22]. As such, the aggregates are not bound to the hydrogel and can pull out of or slide into the lumen easily, which can disrupt the consistency of the cytoarchitecture. We have also observed no significant degradation of agarose columns 3 months after implantation. It is important to consider a bioactive hydrogel with expansive opportunities to tune specific properties and functionalities.

**Fig. 1.**
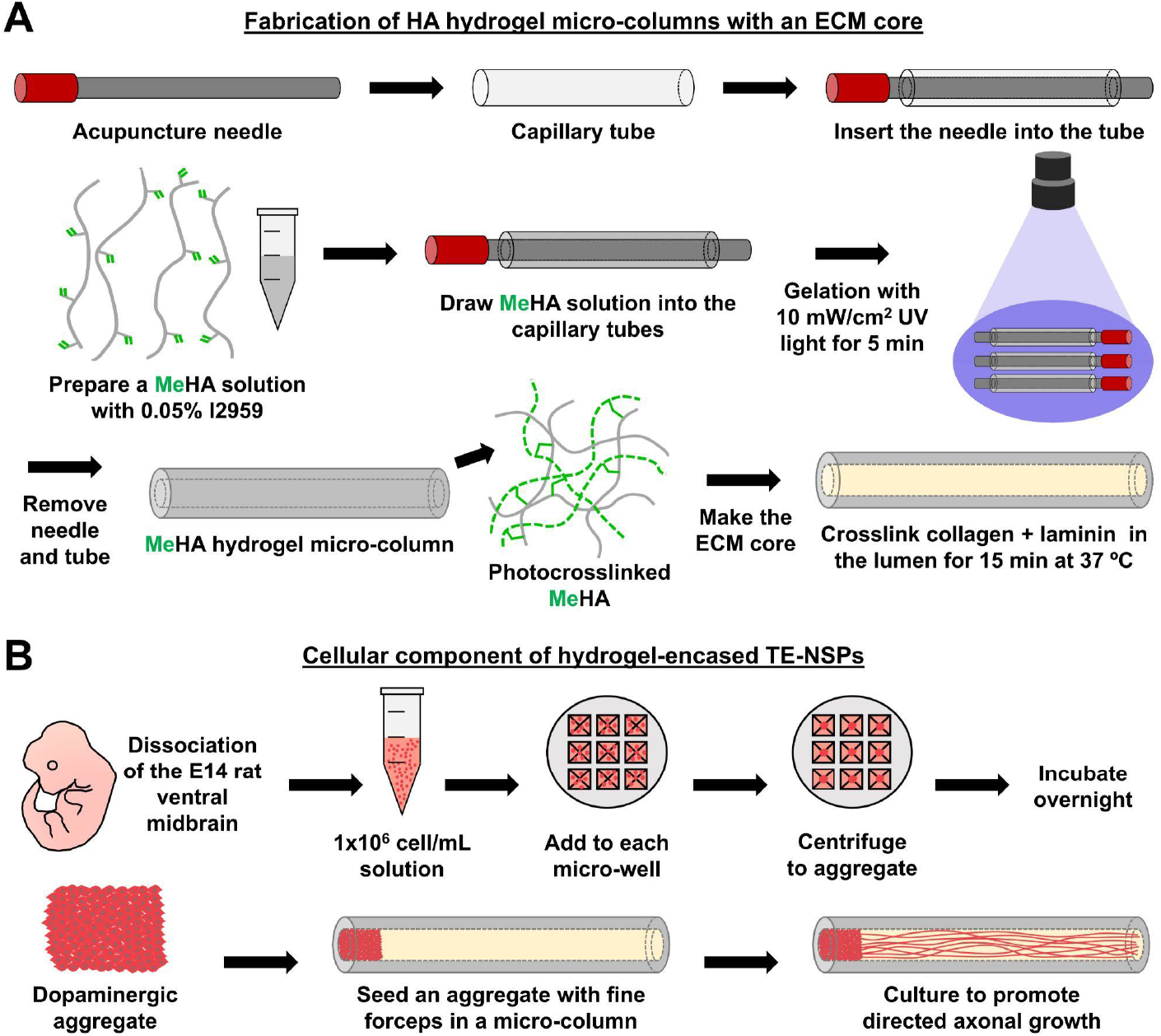
Fabrication methods for the hyaluronic acid-encased tissue-engineered nigrostriatal pathway (TE-NSP). (**A**) The hydrogel encasement for TE-NSPs was made by drawing a MeHA/photoinitiator solution inside a capillary tube containing an acupuncture needle. The schematic shows the modified HA chains, with the reactive methacrylate groups colored in green. Then, the MeHA/photoinitiator solution inside the tubes was crosslinked via exposure to UV light, which permits kinetic chain growth through the reaction of pendant methacrylates. The needle and tube were then removed to obtain the MeHA hydrogel micro-column, with its outer and inner diameter controlled by the choice of tube and needle, respectively. Finally, the ECM core was made by crosslinking a collagen/laminin solution within the lumen. (**B**) In parallel, dopaminergic neurons were obtained from the ventral midbrain of E14 rat pups. A chosen volume of dissociated cell solution was added to micro-wells, centrifuged, and incubated overnight to form an aggregate of neurons. The previous steps converged with the seeding of an aggregate inside one end of the ECM-filled core of the micro-columns using fine forceps. The constructs were then cultured for the desired time to promote axon growth.

In this work, we aimed to advance TE-NSPs by fabricating these constructs with a hyaluronic acid (HA) hydrogel. HA is a major organizing component of the brain ECM and a polysaccharide that consists of repeating D-glucuronic acid and *N*-acetyl-D-glucosamine disaccharide units [23,24]. HA hydrogels have highly controllable properties and can be readily modified with pendant functional groups to confer specific functionalities [14,23,24]. HA has been used extensively for medical applications [23,24] and studied for cell encapsulation for stroke [14] and PD [16], where it has improved the post-transplantation survival of delivered cells. When compared to the inert agarose encasement, HA hydrogels are bioactive and offer improved tunability, allowing their material properties to be modulated according to specific design criteria and applications. Here we focused on methacrylated HA (MeHA), which may be crosslinked via free radical photocrosslinking to form hydrogels [24]. The properties of MeHA hydrogels may be tuned by adjusting the molecular weight of HA, the degree of methacrylation, the macromer concentration, and/or the crosslinking conditions [23]. In this manuscript, we report on the use of MeHA to fabricate TE-NSPs that resemble the nigrostriatal pathway and that feature robust neurite growth, the proper axonal cytoarchitecture, and evoked dopamine release. We also characterize MeHA-encased TE-NSPs after transplantation, by studying the host inflammatory response to the biomaterial and their acute survival.

## Materials and Methods

All procedures were approved by the Institutional Animal Care and Use Committees at the University of Pennsylvania and the Michael J. Crescenz Veterans Affairs Medical Center and adhered to the guidelines set forth in the NIH Public Health Service Policy on Humane Care and Use of Laboratory Animals (2015).

### MeHA synthesis and characterization

As previously reported [25], MeHA was synthesized by the esterification of hyaluronic acid (Lifecore Biomedical, ~60 kDa) with methacrylic anhydride (Sigma Aldrich, 276685) in deionized water for ~3.5 hr at pH 8.5. The reaction was purified by dialysis for 5-7 days, and the product was recovered by lyophilization. The degree of methacrylation (~40%) was determined via ^1^H-NMR by integration of the vinyl singlets (δ~5.8 and δ~6.3 ppm, 1H each) relative to the sugar ring of HA (10H). To study gelation kinetics, 3% and 5% weight/volume (w/v) MeHA solutions were prepared in Dulbecco’s phosphate buffered saline (DPBS; ThermoFisher, 14190136) with 0.05% w/v Irgacure 2959 (I2959; Sigma Aldrich, 410896) photoinitiator. The solution was transferred between the fixed plate and the geometry (20 mm diameter acrylic cone and plate geometry, 59 min 42 s cone angle, 27 μm gap) of a stress-controlled rheometer (TA Instruments, AR2000) fitted to an ultraviolet (UV) lamp (Excelitas Technologies, OmniCure S1500, 320-390 nm). Time sweeps (0.5% strain, 1 Hz) were performed and the storage and loss moduli were recorded before and after exposure to UV light (at *t* = 2 min, *I* = 10 mW/cm^2^) to observe crosslinking *in situ*. The data presented here are representative examples of at least duplicate profiles for each hydrogel formulation. Hydrogels were also cast in ~4.78 mm molds (*n* = 5), and their compressive moduli were measured via dynamic mechanical analysis (TA Instruments, DMA Q800). Specifically, the modulus was calculated as the slope in the 10-20% strain region from generated stress vs. strain curves (0.2 N/min loading rate).

### Fabrication of hydrogel micro-columns

MeHA micro-columns were created by drawing up MeHA solution (with 0.05% I2959 in DPBS) by capillary action into glass capillary tubes (inner diameter: 345 or 398 μm; Drummond Scientific, 10000030, 10000040) containing an inserted acupuncture needle (outer diameter: 160 μm; Lhasa OMS, SJ.16X30). The needle created the inner diameter (ID) and lumen of the micro-column, while the capillary tube formed the outer diameter (OD) and hydrogel shell. The solution was then crosslinked with UV light (*I* = 10 mW/cm^2^) for 5 min. The needles were manually pulled from the tubes, leaving the gelled micro-columns inside. The hydrogels were pushed out of the tubes and into DPBS using 30 gauge needles (BD, 305128). The micro-columns were sterilized with UV light for 30 min and cut to the desired length. For agarose micro-columns, the same procedure was followed, except that 1% w/v agarose solutions were made by heating and stirring agarose (Sigma Aldrich, A9539) in DPBS and then gelled by cooling. The final step before cell seeding was filling the lumen with ECM, which was prepared with 1 mg/mL rat tail type I collagen (Corning, 354236) and 1 mg/mL mouse laminin (Corning, 354232) in Neurobasal (ThermoFisher, 21103049) at 7.2-7.4 pH. After aspirating the DPBS from the surroundings and interior of each micro-column, ECM solution was added into the lumen in excess and crosslinked for 15 min at 37 °C. Fresh media was added to the columns, the excess ECM adhered to the sides of the columns was removed with forceps, and the micro-columns were incubated at 37 °C until seeding **(Fig. 1A)**.

### Neuronal cell culture, aggregation, and seeding

Rat dopaminergic neurons were isolated from embryonic day 14 (E14) rat pups as previously published [26]. Pregnant Sprague Dawley rats (Charles River) were euthanized with carbon dioxide and decapitation, the pups were extracted, and the ventral midbrain was dissected from the brains in Hank’s balanced salt solution (HBSS; ThermoFisher, 14175079). The tissue was rinsed, dissociated with accutase (ThermoFisher, A1110501) for 10 min at 37 °C, and triturated with a pipette. After centrifugation at 200 g for 5 min, the cells were resuspended at 1-2×10^6^ cells/mL using media with Neurobasal, 2% B27 (Invitrogen, 12587010), 1% fetal bovine serum (FBS; Sigma, F0926), 2 mM Glutamax (ThermoFisher, 35050061), 0.3% penicillin-streptomycin (ThermoFisher, 15140122), 4 ng/mL mouse recombinant basic fibroblast growth factor (bFGF; Fisher PMG0034), and 100 μM ascorbic acid (Sigma, A5960). For micro-column seeding, dissociated dopaminergic neurons were first aggregated by adding 15 μL to each micro-well in a custom-made array of 3 × 3 inverted pyramidal micro-wells that fit in 12-well plates. The cells formed aggregates at the bottom of the micro-wells after centrifugation at 1,500 rpm for 5 min, and were subsequently incubated overnight. The aggregates were then pipetted out and inserted with fine forceps to fit within one end of the lumen of the micro-columns **(Fig. 1B)**. After seeding, the micro-columns had approximately 3,000 cells per aggregate. These seeded micro-columns were cultured at 37 °C and 5% CO_2_, with media changes every 2-3 days. In some cases, the rat ventral midbrain neurons were virally transduced to express green fluorescent protein (GFP). In these cases, rat TE-NSPs were incubated overnight at 5 DIV with media including 1/2000 of pAAV1.hSyn.eGFP.WPRE.bHG (final titer of ~3×10^10^ genomic copies/mL; Addgene, 105539-AAV1), with a full media change on the next day.

### Growth characterization of TE-NSPs

The effect of biomaterial encasement on neurite growth length and rate in rat TE-NSPs (OD: 398 μm; ID: 160 μm; length: ~5 mm) was evaluated at 1, 3, 7, and 10 DIV for 1% agarose (*n* = 7), 3% MeHA (*n* = 13), and 5% MeHA (*n* = 14). The lengths were measured from phase contrast images as the distance between the inner edge of the aggregate and the tip of the longest observable neurite using Fiji/ImageJ [27]. The growth rates were estimated with the difference between the neurite length at the current and previous time points divided by the difference in time. The images were obtained using a Nikon Eclipse Ti-S microscope and QiClick camera integrated with NIS Elements software (Nikon). Where applicable, the results were analyzed using repeated measures two-way ANOVA and post-hoc Tukey’s multiple comparison test for differences between groups, with *p* < 0.05 for statistical significance.

### Phenotypic and structural characterization of TE-NSPs

The cytoarchitecture of TE-NSPs and the presence of neuronal and dopaminergic phenotypes were verified by immunolabeling and confocal imaging. TE-NSPs were fixed at 14 DIV in 4% paraformaldehyde (Electron Microscopy Sciences, 15710) in DPBS for 35 min. Whole constructs were permeabilized for 1 hr with 0.3% Triton X-100 (Sigma Aldrich, T8787) in 4% horse serum (ThermoFisher, 16050122) and then incubated overnight with primary antibodies in 4% horse serum at 4 °C. The primary antibodies used in this study included tyrosine hydroxylase (TH; 1/500, sheep, Abcam, ab113), the enzyme in the rate-limiting reaction of dopamine synthesis, and β-tubulin III (1/500, mouse, Sigma, T8578), a neuronal microtubule protein. Then, the cultures were exposed to appropriate fluorescent secondary antibodies (1/500, ThermoFisher, Alexa-488, Alexa-568, Alexa-647) in 4% horse serum for 2 hr at room temperature. Finally, the constructs were incubated for 10 min with Hoechst (1/10000, ThermoFisher, 33342) to stain the nuclei, and rinsed thoroughly. Immunolabeled cultures were imaged with a Nikon A1RSI laser scanning confocal microscope. All images included in this manuscript represent the maximum intensity projection of the full-thickness z-stacks.

### Functional characterization with fast scan cyclic voltammetry

TE-NSPs were assessed for the effect of biomaterial encasement on functionality by measuring electrically-evoked dopamine release with fast scan cyclic voltammetry. TE-NSPs encased in 1% agarose (*n* = 5), 3% MeHA (*n* = 4), and 5% MeHA (*n* = 5) were evaluated at 36-37 DIV. These constructs were incubated with 100 μM L-DOPA (Sigma Aldrich, D9628) for 30 min before transferring to the recording chamber. Here, individual TE-NSPs were perfused at 37 °C with media bubbled with 95% O_2_ and 5% CO_2_, and containing Neurobasal, 2.0 mM L-glutamine, 100 μM ascorbic acid, and 100 μM L-DOPA. A bipolar stimulating electrode (Plastics One) was placed to span a region of interest in the dopaminergic aggregate, and a carbon fiber electrode (150-200 μm outer fiber length, 7 μm diameter) was located in this same area. The stimulation electrode was set to apply a monophasic+ electrical train of 10 pulses of 5 ms width at 20 Hz and a current of approximately 650-750 μA. For recording, the potential of the carbon fiber electrode was scanned linearly from −0.4 V to 1.2 V to −0.4 V vs. Ag/AgCl at a rate of 400 V/s with a voltammeter/amperemeter (Chem-Clamp; Dagan Corporation). The stimulation train was applied at 8 s, and cyclic voltammograms (CVs) were recorded every 100 ms during the entire session using the Demon Voltammetry and Analysis Software [28]. The current at the peak oxidation potential of dopamine in consecutive CVs and all the CVs obtained during the session for one construct were averaged from 4-6 recording runs within the same location 8-12 min apart. This averaging was done to minimize the noise and emphasize the signal. The currents were converted to dopamine concentration with a calibration curve. We recorded the elicited current after injection of 1.5, 3, and 6 μM dopamine hydrochloride (Sigma Aldrich, H8502) and obtained the slope of the regression of concentration vs. current. The dopamine in this case was dissolved in deionized water with 126 mM NaCl, 2.5 mM KCl, 1.2 mM NaH_2_PO_4_-H_2_O, 2.4 mM CaCl_2_-2H_2_O, 1.2 mM MgCl_2_-6H_2_O, 25 mM NaHCO_3_, and 0.4 mM L-ascorbic acid (all from Sigma Aldrich). Dopamine concentration peaks after release were analyzed using the Kruskal-Wallis test with Dunn’s multiple comparisons test. Each dopamine concentration that was analyzed corresponded to one construct and was obtained from the average of several stimulation/recording runs as explained above.

### Transplantation of TE-NSPs

Male Sprague Dawley rats (8-11 weeks, 275-290 g; Charles River) were used for proof-of-concept transplants of MeHA-encased TE-NSPs at 14 DIV, with a terminal time point of 2 weeks post-implantation (3% MeHA: *n* = 3; 5% MeHA: *n* = 3). Male athymic rats (12-13 weeks, 260-280 g; Charles River) were used for implants of acellular micro-columns with a terminal time point of 6 weeks (1% agarose: *n* = 3; 3% MeHA: *n* = 5; 5% MeHA: *n* = 5). Athymic rats were used as a transition to future studies using athymic rats lesioned to have parkinsonian symptoms. Rats were anesthetized with 5% isoflurane (Midwest Veterinary Supply, 193.33165.3) and mounted in a stereotactic frame, with anesthesia being maintained at 2.5%. The scalp was disinfected with betadine, and 2.0 mg/kg bupivacaine (Hospira, NDC 0409-1161-19) was injected in the incision area. The incision was made along the midline of the scalp, and a 5 mm round craniectomy was made at +5 mm AP and 2.3 mm lateral relative to Bregma. TE-NSPs were pulled into a needle (Vita Needle; OD: 534 μm, ID: 420 μm) containing a rod that lay directly in contact with the construct. A protective polyurethane tubing (OD: 686 μm; ID: 559 μm; Microspec Corporation) was cut to fit the needle, sealed on one end, and used to cover the outer area of the needle. The needle was inserted into a Hamilton syringe attached to a stereotactic arm, and its plunger was put directly in contact with the exposed rod coming out of the needle. The stereotactic arm was adjusted to be at 38° relative to the horizontal, the dura was opened, and the needle was lowered ~12 mm into the brain. Then, the sheath was pulled up with forceps to break the seal, and the TE-NSPs were laid out by pulling up the needle ~5 mm while a stationary arm maintained the plunger in place. After 2 min, the needle was pulled out entirely (with the plunger facing no resistance), the scalp was sutured, and the animals received a subcutaneous injection of 0.05 mg/kg buprenorphine (Reckitt Benckiser, NDC 12496-0757-1). At the terminal time points, the animals were anesthetized with isoflurane and sacrificed via transcardial perfusion with heparinized saline followed by 10% formalin. The brains were extracted, post-fixed overnight in 10% formalin, and stored in 1X PBS at 4 °C.

### Characterization of the host brain response

The brains of animals implanted with acellular micro-columns were placed in a custom-made mold with indentations at an angle orthogonal to the implant trajectory, cut into 2 mm thick blocks containing axial cross-sections of the micro-columns, and processed through paraffin. Eight micron-thick sections were collected on Superfrost Plus slides (Fisher Scientific, 12-550-15). These sections were exposed to two washes with xylene (Fisher Scientific, X5-500) for 8 min each, two washes with 100% ethanol (Leica Biosystems, 3803686) for 1 min each, two washes with 95% ethanol (Leica Biosystems, 3803682) for 1 min each, and a wash with deionized water. Following the dewaxing stage, antigen retrieval was completed in Tris EDTA buffer pH 8.0 using a microwave pressure cooker. After returning to room temperature and rinsing with water, each section was blocked for 30 min with normal horse serum (Vectastain Elite Universal Kit; Vector Laboratories, PK-6200) dissolved in 1X Optimax Wash Buffer (pH 7.4; BioGenex, HK583-5K). The sections were then incubated overnight at 4 °C with a primary antibody solution made in 1X Optimax buffer. For these studies, the primary antibodies targeted: (1) glial fibrillary acidic protein (GFAP; 1/1000, goat, Abcam, ab53554), the key intermediate filament protein in astrocytes; (2) ionized calcium-binding adapter molecule 1 (IBA1; 1/1000, rabbit, Wako, 019-19741), a calcium-binding protein in microglia, glial cells in the brain involved in its immune response; and (3) neuronal nuclei (NeuN; 1/1000, chicken, Millipore, ABN91), a neuron-specific protein in nuclei. Afterwards, the sections were rinsed three times with 1X Tween/PBS with shaking for 8 min and incubated for 1 hr at room temperature with the secondary antibody solution made in horse serum in 1X Optimax. The secondaries consisted of donkey anti-goat 568, donkey anti-chicken 488, and donkey anti-rabbit 647 (1/1000, Invitrogen). The sections were washed twice with 1X Tween/PBS for 5 min, covered with Fluoromount-G (Southern Biotech, 0100-01) and a coverslip, and imaged with the confocal microscope.

We used the NeuN, GFAP, and IBA1 images to quantify the host response in four layers surrounding the implant cross-section, each located at 50, 100, 150, and 200 μm from the edge of the implant. In all cases, the outcome measures were normalized relative to the contralateral side by locating an area within it at the same coordinates as the implant, and measuring those outcomes in the same four layers. This step was undertaken to isolate the effect of the biomaterial on the response from inherent differences between sections. To quantify host neuron survival around the micro-column, the number of NeuN+ cells was counted within each layer and normalized. To assess the surrounding microglia and astrocyte presence, we calculated and normalized mean intensity of the IBA1 and GFAP signal in each layer, respectively, which corresponded to the total value of the pixels (sum intensity) per area. The creation of the layers and the measurements were carried out using ImageJ/Fiji. Two-way ANOVA was used to determine the significance of the effect of the biomaterial and the layers, while Tukey’s multiple comparisons tests were employed to compare different biomaterials in the same layers.

### Optical clearing and immunolabeling

Thick brain sections were subjected to a clearing and immunohistochemistry protocol to visualize the entire constructs and to assess their cytoarchitecture and survival [29]. A vibratome (Leica, VT1000S) was utilized to obtain 2 mm thick sagittal sections containing the implanted constructs. The sections were dehydrated with increasing concentrations of methanol (Fisher Scientific, A412) and then incubated overnight at room temperature with 66% dichloromethane (Sigma Aldrich, 270997) and 33% methanol for delipidation and refractive index matching. Afterwards, the sections were bleached overnight in chilled 5% H_2_O_2_ (Sigma, H1009), rehydrated with decreasing concentrations of methanol, permeabilized for 2 days at 37 °C, and blocked for 2 days at 37 °C. Permeabilization was performed using 0.16% Triton X-100, 20% dimethyl sulfoxide (DMSO; Sigma Aldrich, 276855), and 0.3 M glycine (Bio-Rad, 161-0718) in 1X PBS, while the blocking solution consisted of 0.16% Triton X-100, 6% horse serum, and 10% DMSO in 1X PBS. Afterwards, the sections were exposed to primary and secondary antibodies for 7 days at 37 °C each, with washing steps in between. The primary antibody buffer was 3% horse serum and 5% DMSO dissolved in 1X PBS with 0.2% Tween-20 (Sigma Aldrich, P2287) and 10 μg/mL heparin (Sagent Pharmaceuticals, 25021-400-10). The secondary buffer contained 3% horse serum and the same solvent. We used primary antibodies for TH (1/100, sheep, Abcam, ab113) and GFP (1/100, mouse, Aves, GFP-1020) and the appropriate Alexa-Fluor secondaries (1/250). The sections were dehydrated with increasing concentrations of methanol, exposed to Visikol® HISTO-1™ overnight at room temperature, and incubated with Visikol® HISTO-2™ at least for 2 hr before imaging with the confocal microscope.

## Results

### Characterization of the mechanical properties of MeHA and agarose hydrogels

We have traditionally fabricated TE-NSPs with 1% agarose hydrogels [17], but here we focused on HA hydrogels, where ~40% of the primary alcohol groups in the *N*-acetyl-D-glucosamine units of HA were modified with methacrylates to create MeHA. MeHA forms hydrogels via free radical photocrosslinking, where light first generates radical species via the photolysis of a photoinitiator, and these free radicals then propagate via methacrylates to form kinetic chains [30]. Hydrogels were created by irradiating MeHA/photoinitiator solutions with UV light, as confirmed with rheological measurements showing an increase in the storage (G’) modulus as a function of time after exposure to light to values much greater than the loss (G”) modulus **(Fig. 2A)**. As expected, the G’ value for a 5% MeHA hydrogel was greater than that for 3% MeHA. Moreover, the profiles demonstrated that a light exposure time of ~5 min was sufficient to reach a plateau in moduli. We further investigated how the mechanical properties of MeHA and agarose hydrogels compare by obtaining the compressive moduli from stress vs. strain curves **(Fig. 2B)**. The compressive modulus for 5% MeHA was significantly greater than the value for 3% MeHA, and both MeHA hydrogels were significantly stiffer than 1% agarose **(Fig. 2C)**.

**Fig. 2.**
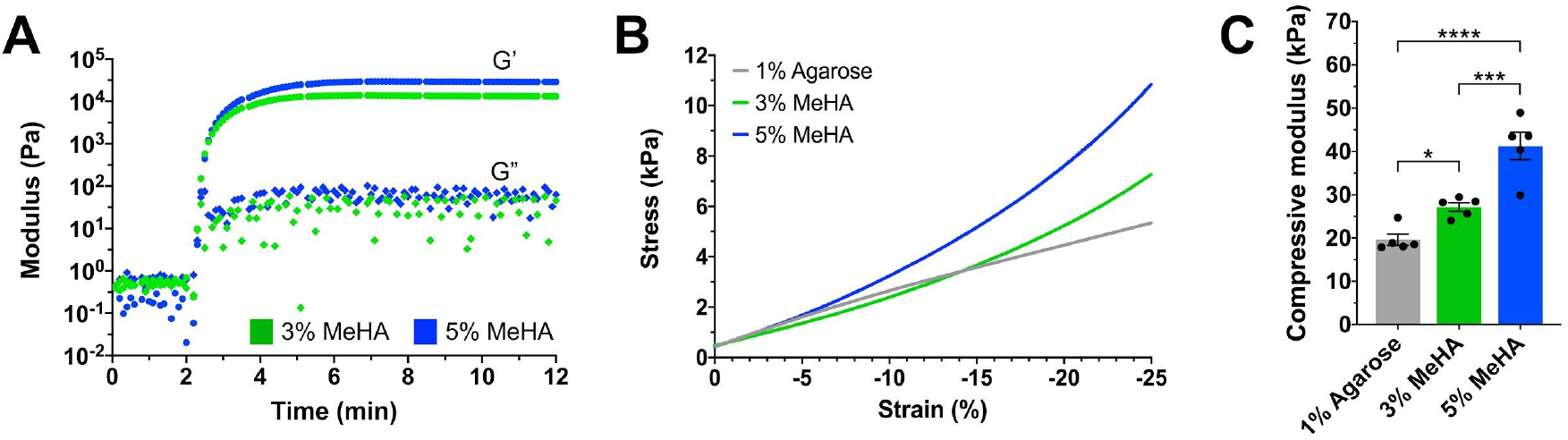
Mechanical assessment of MeHA and agarose hydrogels. (**A**) Photorheology profile shows the change in the storage (G’) and loss (G”) moduli of 3% and 5% MeHA solutions after exposure to UV light at 2 min. (**B, C**) The compressive moduli were calculated from the slope of the stress vs compressive strain curves. Data presented as mean ± SEM (\**p* < 0.05, \*\*\**p* < 0.001, \*\*\*\**p* < 0.0001).

### MeHA is a suitable encasement for TE-NSPs based on growth outcomes

Initially, we confirmed that MeHA hydrogels could be produced in the shape of a hollow micro-column and filled with crosslinked collagen and laminin. Qualitatively, we have observed that the lumen of MeHA micro-columns can be completely filled with ECM, both axially and longitudinally, with more consistency than agarose. We produced MeHA-encased TE-NSPs with rat embryonic dopaminergic neurons and ~0.5 cm long micro-columns and tested several outcome measures *in vitro*. Biomaterial type and culture duration had significant effects on both neurite growth lengths and rates. All mean lengths were higher for MeHA groups relative to agarose, with significant differences being observed between 1% agarose and at least one of the MeHA groups at all time points **(Fig. 3A)**. In many cases 5% MeHA performed better than 3% MeHA in terms of mean length. The greatest growth rates were observed in the 3-7 DIV range for all encasements, and the lowest were seen at 1 and 10 DIV **(Fig. 3B)**. The axons extended completely through the lumen by 10 DIV, which may explain the decreased growth rates as the neurites reached the end of the biomaterial. The mean growth rates were generally higher for MeHA in all cases, and there were significant differences between 1% agarose and MeHA groups at 1, 3, and 7 DIV. In one case, the growth rate in 5% MeHA was significantly higher than 3% MeHA at 7 DIV. MeHA constructs exhibited the correct cytoarchitecture and phenotypic expression, including segregated neuronal somata and dopaminergic axon tracts, similar to the nigrostriatal pathway, as noted from Hoechst+ nuclei and TH+ neurites, respectively **(Fig. 3C–F)**. While we observed instances of neuronal migration/dispersal along the axon tracts, the presence of somata in the axon region was minimal.

**Fig. 3.**
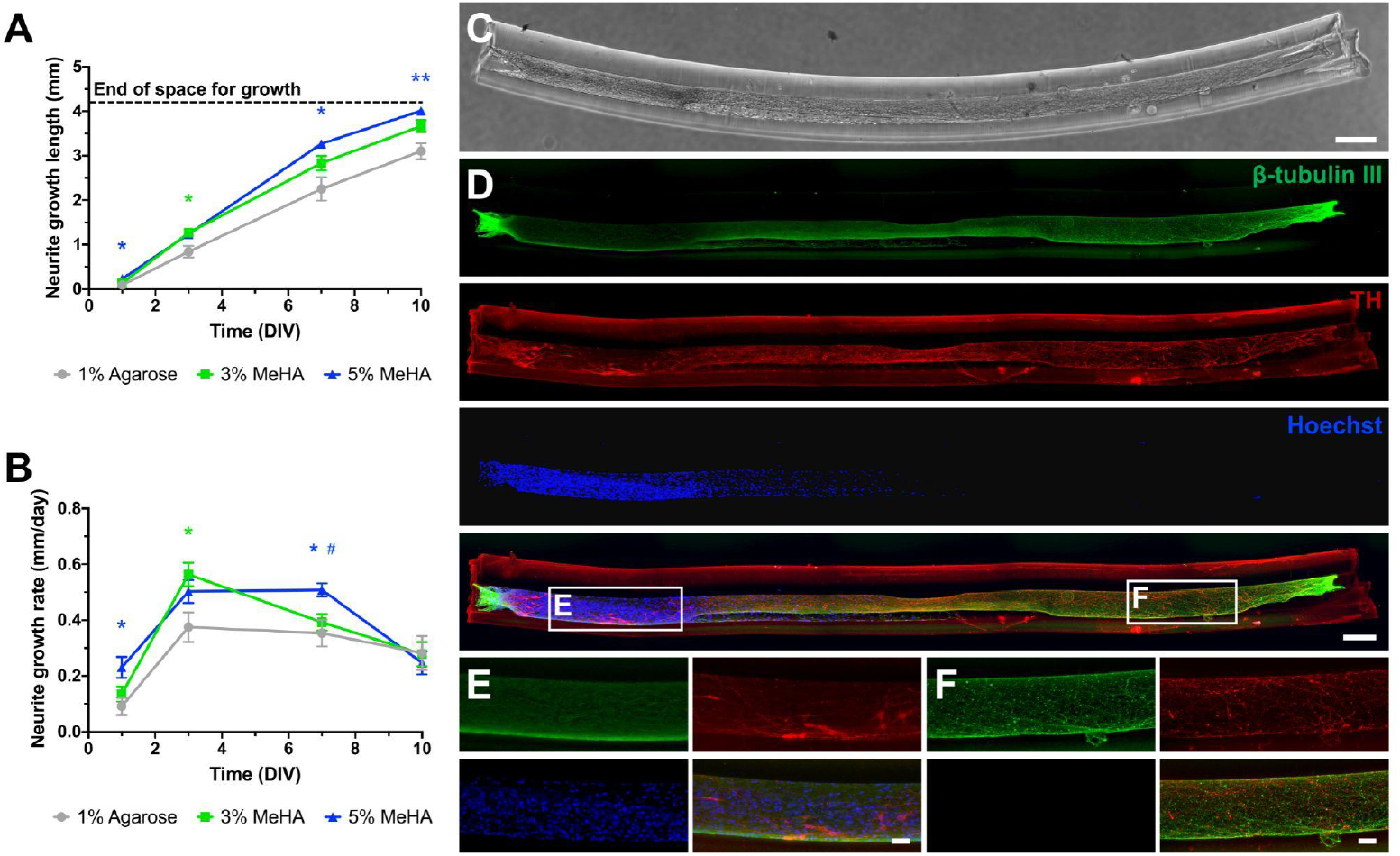
Growth, phenotype, and cytoarchitecture of MeHA-encased TE-NSPs. (**A, B**) The growth lengths and rates for neurites extended by aggregates of rat embryonic ventral midbrain neurons were quantified as a function of time in 1% agarose (*n* = 7), 3% MeHA (*n* = 13), or 5% MeHA (*n* = 14) micro-columns (OD: 398 μm, ID: 160 μm, length: ~0.5 cm). Repeated measures two-way ANOVA yielded a significant effect for the biomaterial over length (*p* = 0.0002) and rate (*p* = 0.0011) and for time over length and rate (*p* < 0.0001). Data presented as mean ± SEM (\**p* < 0.05, \*\**p* < 0.01 from Tukey’s multiple comparisons test; green and blue asterisks indicate significant differences for 3% and 5% MeHA relative to 1% agarose, respectively; # represents the comparison between 5% and 3% MeHA with *p* < 0.05). (**C**) Phase-contrast image of a representative 3% MeHA TE-NSP at 14 DIV. (**D**) Maximum intensity projection of confocal z-stacks after staining the construct in **C** for axons (β-tubulin III; green), dopaminergic neurons (tyrosine hydroxylase (TH); red), and nuclei (Hoechst; blue). (**E, F**) Higher magnification images of the aggregate and axon tract regions, respectively. Scale bars: (C) 250 μm; (D) 200 μm; (E, F) 50 μm.

### MeHA is a suitable encasement for TE-NSPs based on functional outcomes

Following these characterization studies, we examined the functionality of TE-NSPs encased in MeHA in terms of electrically-evoked dopamine release. We used fast-scan cyclic voltammetry (FSCV), a technique with high temporal/spatial resolution used to measure changes in neurotransmitter concentration based on the currents elicited by their oxidation or reduction [31]. We introduced a carbon fiber electrode into the aggregates to record extracellular dopamine release after electrical stimulation **(Fig. 4A)**. The evoked dopamine concentration for one construct was calculated from one location within the aggregate after averaging the current traces and cyclic voltammograms (CVs) from several recording sessions. Rat TE-NSPs in 3% and 5% MeHA had a mean dopamine release of 317.6 ± 115.7 nM and 206.289 ± 35.69 nM, respectively, in comparison to 273.1 ± 45.23 nM when encased in 1% agarose, which did not reflect a significant effect of the biomaterial used on functionality **(Fig. 4B)**. In all three encasement types, the cells responded to stimulation **(Fig. 4C–E, top),** and the released molecules exhibited the chemical signature of dopamine. The CVs at peak release **(Fig. 4C–E, center)** showed clear current peaks at the oxidation potential and troughs at the reduction potential of dopamine around 0.6 V and −0.3 V, respectively. CVs recorded as a function of time also show that characteristic dopamine signal throughout the duration of release **(Fig. 4C–E, bottom)**.

**Fig. 4.**
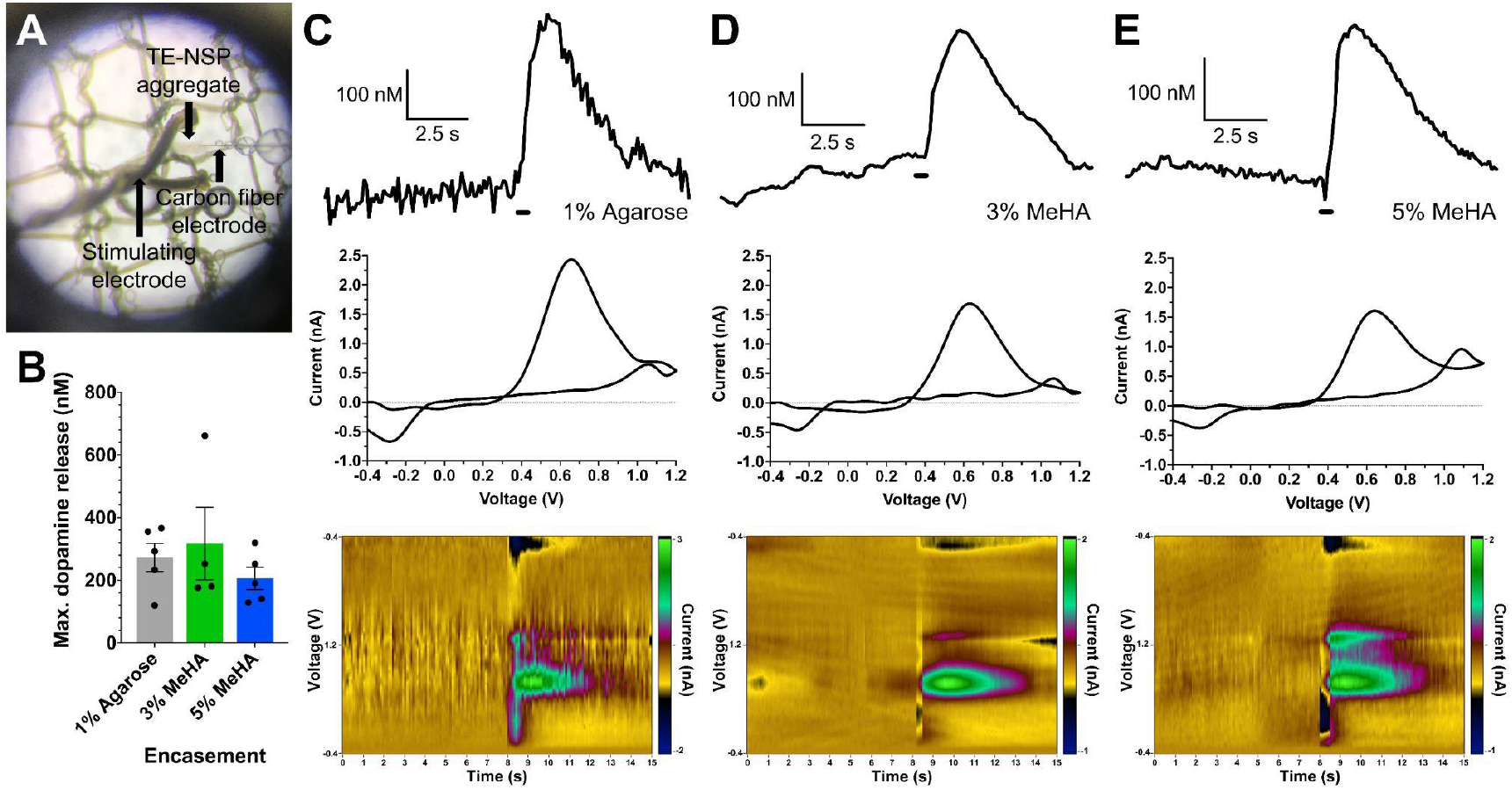
Dopamine release functionality of MeHA-encased TE-NSPs. (**A**) At 36-37 DIV, TE-NSPs were analyzed with FSCV for electrically-evoked dopamine release. A carbon fiber electrode was inserted into the aggregate, and a stimulation electrode was placed to span it. (**B**) The effect of biomaterial encasement on dopamine release was evaluated, with no significant differences (mean ± SEM, *p* = 0.6175). Each dot represents the evoked dopamine concentration from one construct, averaged from several stimulation/recording sessions. (**C-E**) Concentration traces (top), cyclic voltammograms (CV) at peak release (center), and CV colorplots (bottom) for constructs in 1% agarose and 3% and 5% MeHA. The black bar represents the time of stimulation. The plots indicate the release of dopamine, as assessed from characteristic oxidation and reduction currents around 0.6 V and −0.3 V, respectively.

### MeHA promotes a less intense host response after implantation along the nigrostriatal pathway

Having confirmed that MeHA can be used to create TE-NSPs with the intended structure and function, we proceeded to assess these constructs *in vivo*. We sought to determine the host response, in terms of NeuN+ neuron, IBA1+ microglia, and GFAP+ reactive astrocyte presence, around implanted MeHA micro-columns and to compare this to 1% agarose **(Fig. 5A–C)**. Within a layer 50 μm from the edge of the implant, we observed that host NeuN+ neurons surrounding it decreased to 20-53% of the number in the contralateral side **(Fig. 5D, left)**. Farther away from the edge, the normalized NeuN+ counts approached 1, signifying that the effect is distance-dependent and that host neurons are better protected farther from the implants. Indeed, two-way ANOVA indicated that the layer distance had a significant effect on normalized cell counts. The biomaterial type approached significance but did not meet it (*p* = 0.0887). Still, the data show that within 50 μm, on average more host neurons survived with MeHA implants (46 and 53% for 3% and 5% MeHA, respectively) when compared to agarose (20%). Moreover, host neurons around 5% MeHA exhibited survival comparable to the contralateral side faster than the other groups, as the normalized count quickly approached 1 after the second layer (50-100 μm from the edge) and beyond, while the others exhibited the same only at the farthest layer.

**Fig. 5.**
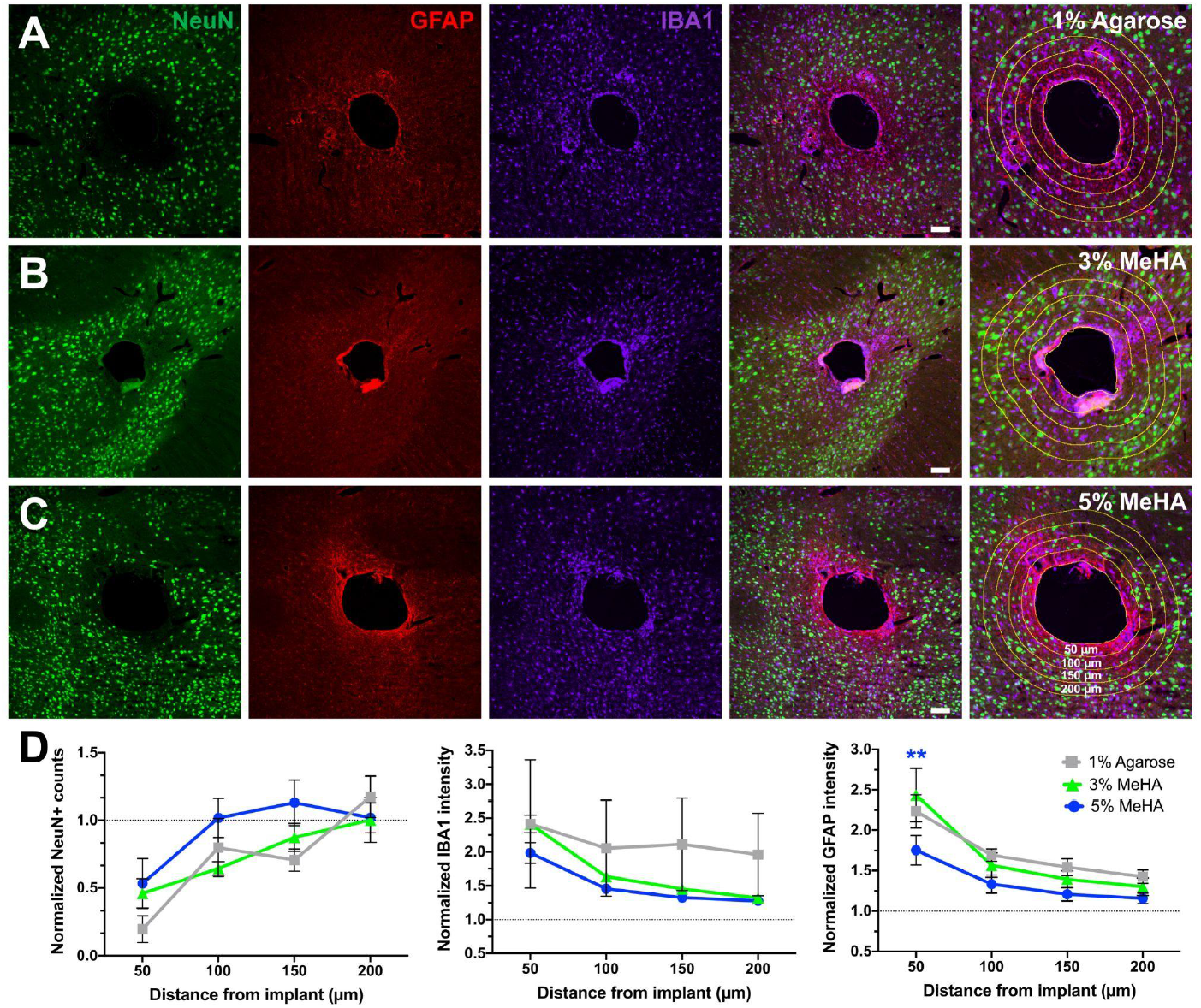
Rat brain response to the implantation of MeHA and agarose hydrogel columns along the nigrostriatal pathway. **(A-C)** Acellular hydrogel micro-columns (OD: 345 μm, ID: 160 μm, length: ~0.5 cm) fabricated with 1% agarose (*n* = 3), 3% MeHA (*n* = 5), and 5% MeHA (*n* = 5), respectively, were implanted along the nigrostriatal pathway of athymic rats for 6 weeks to assess the host response. Sections orthogonal to the implant trajectory were stained for neurons (NeuN; green), astrocytes (GFAP; red), and microglia (IBA1; far red) to determine the extent of host cell death and the inflammatory response around the micro-columns. **(D)** The number of NeuN+ cells and the staining intensity of IBA1+ and GFAP+ cells in the injection side were normalized to the contralateral side and quantified in four distinct layers located at 0-50, 50-100, 100-150, and 150-200 μm from the edge of the implant, as observed in the far-right images in **A**-**C**. Two-way ANOVA tests indicated a significant effect for layer distance in the case of NeuN+ counts (*p* < 0.0001) and for layer distance and biomaterial type in the case of IBA1 (*p* = 0.0183, *p* = 0.0256) and GFAP intensity (*p* < 0.0001, *p* = 0.0059), respectively. Data presented as mean ± SEM (\*\**p* < 0.01 between 3% and 5% MeHA for GFAP at the 50 μm layer). Scale bars: 100 μm.

In the case of the host inflammatory response, we noticed that both IBA1+ microglia and GFAP+ astrocyte presence was approximately twice as high as the contralateral side in the layer closest to the implant, and their intensity decreased to values closer to the contralateral side farther away **(Fig. 5D, center, right)**. In general, the host tissue had a lower normalized microglia and astrocyte intensity (*i.e.*, closer to contralateral values) around both types of MeHA micro-columns when compared to 1% agarose at the same layer distance. For example, in the second layer (50-100 μm from the implant) the mean normalized IBA intensity values were 1.64 ± 0.08 and 1.46 ± 0.07 for 3% and 5% MeHA, respectively, and 2.05 ± 0.71 for 1% agarose; for GFAP the corresponding values were 1.57 ± 0.15, 1.33 ± 0.11, and 1.69 ± 0.08, respectively. Similar gaps between these biomaterials were observed in the other layers. These observations are strengthened by the significant effect of both layer distance and biomaterial type on normalized IBA1 and GFAP intensity. In addition, the average host response was less intense with 5% MeHA. Despite this observation, the only significant difference between groups was seen between 3% and 5% MeHA for GFAP staining in the first (0-50 μm) layer (*p* = 0.0081).

### MeHA-encased TE-NSPs survive after implantation along the nigrostriatal pathway in rats

A crucial consideration for the use of MeHA for TE-NSPs was whether these constructs could maintain their axonal structure during and after implantation and survive over time. After 14 DIV, MeHA-encased TE-NSPs were implanted to span the nigrostriatal pathway in rats **(Fig. 6A)**. MeHA TE-NSPs could be loaded into needles and manipulated without observable damage to the integrity of the cells and axon tracts. Through tissue clearing and immunohistochemistry we observed that the constructs could be delivered to span the nigrostriatal pathway, with one end in the striatum **(Fig. 6B)** and the other end proximal to the nigral cell bodies **(Fig. 6C)**, and maintain their cytoarchitecture and integrity for at least 2 weeks. We also noticed some instances of putative outgrowth from the TE-NSP and ingrowth from host TH+ neurons into one end of the constructs **(Fig. 6B–C)**, although other studies are needed to confirm and elucidate its extent.

**Fig. 6.**
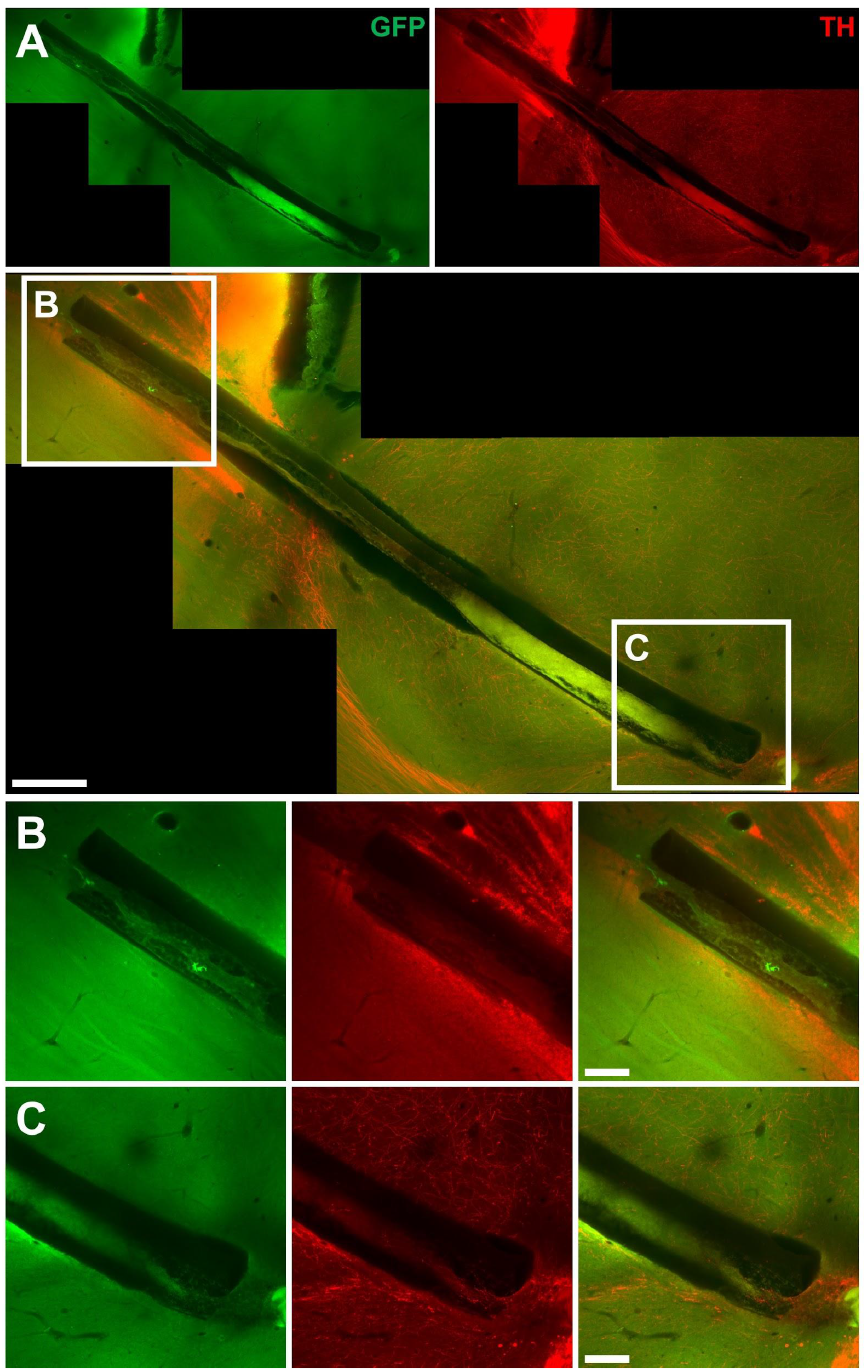
Implantation of MeHA-encased TE-NSPs. (**A**) Confocal image of a GFP+ TE-NSP encased in 3% MeHA (OD: 398 μm, ID: 160 μm, length: 0.6 cm), implanted for 2 weeks in a rat, and stained for all neurons (GFP; green) and dopaminergic neurons (TH; red). (**B, C**) Magnification of areas of the implant within the striatum and adjacent to the nigra, respectively, showing the correct delivery trajectory. Scale bars: (A) 500 μm; (B, C) 200 μm.

## Discussion

Current therapies for treatment of PD center on drug delivery, DBS, or cellular replacement strategies. While these approaches have had varying degrees of success, none address the importance of the anatomical connection between the substantia nigra and striatum. To tackle this, we developed TE-NSPs consisting of an aggregate of dopaminergic neurons and their axon tracts surrounded by an agarose micro-column. In this study we improved upon this design by constructing TE-NSPs with HA (modified as MeHA) as the biomaterial for the encasement. HA is the major component of the brain ECM and can bind to neural cells through cell surface receptors such as CD44 and CD168 [32]. It forms complexes that offer the structural support and modulation of development, cell signaling, intercellular interactions, and synaptic plasticity that are key to brain function and response to injury [33,34]. HA hydrogels have been used for 3D stem cell encapsulation and to control their expansion, migration, and differentiation [35–40]. In neural applications, HA hydrogels have been utilized to differentiate human neural progenitor cells (NPCs) and to augment survival and behavioral outcomes in stroke models [12,39]. Human embryonic stem cell-derived dopaminergic neurons have been encapsulated in HA to mature them and to protect them during transplantation [16]. This biomaterial can form hydrogels when the backbone is modified with crosslinkable functional groups [23,24]. We used MeHA as a start, given its status as a common HA derivative for photocrosslinking. A photocrosslinking paradigm may expand the future capabilities of TE-NSPs, as it enables *in situ* cell encapsulation and permits control over gelation and hydrogel properties [23]. MeHA and other modified HA hydrogels have also been used extensively for 3D bioprinting [41–44], which aligns with our long-term goal of standardizing our micro-column fabrication protocol so that it is highly reproducible and more scalable than our current manual approach.

We initially characterized the growth, structure, and function of MeHA-encased TE-NSPs fabricated with rat cells to ascertain the feasibility of using this hydrogel. MeHA can be applied to build hollow micro-columns containing a lumen of crosslinked collagen and laminin. Neuronal aggregates sourced from rat embryonic ventral midbrains and seeded within these columns were able to extend long axon tracts that stained for TH, a key dopaminergic marker. Growth and functional outcomes were compared to 1% agarose micro-columns given that this biomaterial has been our historical standard when fabricating TE-NSPs. The type of hydrogel had a significant effect on growth length and rate, with MeHA-encased constructs exhibiting greater outcomes relative to agarose. The growth rates increased over time until they decreased when the neurites reached the end of the micro-columns, which may imply that the aggregates need time to adhere and adapt to their environment before extending neurites faster. While MeHA may not improve the dopamine release seen with agarose, it did not negatively affect it, as it supported dopamine release comparable to time-matched constructs encased in agarose. Overall, we validated that MeHA can be used to make TE-NSPs that structurally and functionally resemble the nigrostriatal pathway.

The effects on growth could be attributed to MeHA itself providing a microenvironment more conducive for neurite growth than agarose. For example, HA could be more amenable to cells given its role in the brain ECM. It may be better at locally retaining growth factors in the media in comparison to agarose. On the other hand, the improved growth could be a result of the distinct mechanical properties of MeHA and agarose. We have previously shown that axons have the highest density of growth along the hydrogel-ECM interface in the circumference of the lumen [45]. As such, while the neurites use the ECM core for growth throughout the lumen, they also interact with the hydrogel as a substrate, and its mechanical properties may thus impact cell behavior. Indeed, it has been extensively studied that neural cells and other types respond to substrate stiffness [46–48]. There has been an inverse relationship between neurite growth and substrate stiffness in studies using mouse hippocampal NPCs seeded on 3D hydrogels of HA-pentenoate and poly(ethylene glycol)-bis(thiol) crosslinker [49], mouse ventral midbrain NPCs in MeHA [50], and chick dorsal root ganglia in 3D agarose hydrogels [51]. Human stem cell-derived NPCs exhibited more spreading and attachment in softer acrylated HA hydrogels [52] and greater neurite outgrowth/density and neuronal differentiation in 3D MeHA hydrogels that were less stiff [53]. While our results coincide with stiffness influencing neurite growth, we observed the opposite: the stiffer MeHA micro-columns performed better than the less stiff 1% agarose. This may imply that other properties are the driving force in our case. MeHA itself may provide a better environment than agarose apart from stiffness, or MeHA may support a more structurally sound encasement that better maintains the integrity of the ECM core. On the latter point, it is common to see shrinking, detachment, or inconsistent crosslinking of ECM inside agarose hydrogels, which may disrupt neurite growth and is not as commonly observed with the MeHA hydrogels. Our results may also reflect that we are using a different type of scaffold, one in which the hydrogel substrate is found in the periphery of a 3D micro-column, while previous studies have relied on 2D surfaces or 3D growth in whole hydrogel pieces. Indeed, in a past publication we demonstrated improved neurite health in micro-tissues similar to TE-NSPs when grown in 3-4% agarose micro-columns relative to 1-2% agarose [54]. A limitation of the current study is that we only compared to 1% agarose, and not to stiffer agarose hydrogels, given that our objective was to use our standard encasement as a control. Regardless, we theorize that this direct relationship between stiffness and growth may be a feature of our micro-column scaffolds.

The next step in our characterization consisted of assessing the host response to the biomaterial on its own. The data showed that the implantation of hydrogel micro-columns along the nigrostriatal pathway causes a noticeable effect in the host tissue that is strongest close to the edge of the implant and weakest farther from it. We showed that on average, more host neurons survive and there is a less intense microglia and astrocyte presence around the micro-column when using MeHA relative to 1% agarose. Moreover, the stiffer 5% MeHA seemed to yield the least pronounced host response on average. While the effect of the type of biomaterial was statistically significant on the microglia and astrocyte response, this was not the case for host neuron survival, and there were mostly no significant differences between groups. A previous study that utilized acrylated HA with conjugated RGD peptide and peptide crosslinkers found that stiffer hydrogels with a storage modulus of ~10^3^ Pa elicited significantly more astrocyte presence in the mouse brain, while the microglia response was not affected [39]. In a stroke model, stiffer hydrogels caused more inflammation, but not more astrocytic presence, which was attributed to masking by the overwhelming activation of astrocytes as a result of the stroke. Another study using a soft, acrylated HA hydrogel containing adhesion peptides, heparin, and a peptide crosslinker also found that the implant did not exacerbate the host response apart from the stroke [55]. While our MeHA hydrogel was different in terms of shape, crosslinking method and pendant ligand presentation, we observed the opposite effect, where the stiffer MeHA hydrogels (storage moduli of ~10^4^ Pa) performed better than 1% agarose. We anticipate this is a result of MeHA being more similar to the brain ECM than agarose. In fact, HA has been shown to reduce the number of astrocytes after brain damage [56]. The slightly better performance of 5% MeHA over 3% MeHA may coincide with a recent study on the role of HA in brain injury and neurodegenerative disease, which concluded that a lesser amount of high molecular weight HA is related with greater astrocyte reactivity and inflammation [57]. It will be relevant to explore how the host response differs in brains lesioned to simulate PD.

The final step in our assessment of MeHA was conducting proof-of-concept implantations with cellular constructs. Specifically, we confirmed that these MeHA TE-NSPs could be successfully delivered into rat brains along the nigrostriatal pathway and that the cells could survive and retain their axonal cytoarchitecture acutely. As such, we deemed that MeHA was a feasible encasement for TE-NSPs in terms of structure, function, and transplantation. More detailed *in vivo* studies will shed more light on the ideal HA derivative or MeHA type for TE-NSP implantation based on properties such as degradation, stiffness, and functionality. Various competing considerations will need to be balanced. It may be advisable to have a slow degrading hydrogel encasement to shield the neurons and axon tracts from inflammation or infiltration of other cell types. MeHA is mainly susceptible to enzymatic degradation by hyaluronidases, and it seemed that under our circumstances there was an insufficient amount for substantial degradation to occur on the timescale of these studies. On the other hand, it may be advisable to have a controllable degradation profile in which there is little to no degradation during the *in vitro* culture and early implant periods, with degradation becoming more prominent *in vivo* as time goes on. This would allow TE-NSPs to grow in 3D with the proper cytoarchitecture, have sufficient integrity to be implanted, and integrate with the host tissue acutely. Subsequent degradation may enable a seamless integration of the dopaminergic axons with the surrounding tissue. Another consideration is the inverse relationship between degradation rate and stiffness and how the latter is associated with the inflammatory response. While the mechanical properties of implanted hydrogels should match those of the brain, they must also possess enough structural integrity for the micro-columns to hold their shape before, during, and after implantation. Properties such as degradation and stiffness can be tuned by modulating the crosslinking density of MeHA hydrogels. Greater control over degradation may be achieved through the use of hydrogels that are more susceptible to hydrolysis or cell-mediated degradation. For example, HA hydrogels sensitive to hydrolysis have been created by modifying HA with methacrylates that are bound to lactic acid or caprolactone, both of which undergo ester hydrolysis [23,58]. Di-thiol peptides sensitive to cleavage by matrix metalloproteinases expressed by cells have also been used for crosslinking [59].

Other types of HA or hydrogels could also be used to make TE-NSPs depending on the desired properties (*e.g.*, degradation, mechanics, binding of cell-responsive peptides, release of growth factors). For example, previous studies have characterized HA hydrogels that consist of two distinct, interpenetrating networks: (1) HA modified with β-cyclodextrin and HA modified with adamantane that form supramolecular networks via guest-host assemblies and (2) a secondary MeHA network crosslinked via thiol-Michael addition [25]. The primary network conferred self-healing properties to the gel system, while the secondary one enhanced its toughness. Additional increases in hydrogel toughness could be achieved when the two networks were tethered covalently. Hydrogels such as these could allow TE-NSPs to recover from mechanical deformation without failure and may provide them with greater resilience to handling during culture and implantation. Other publications have capitalized on sequential crosslinking approaches to control mechanical properties and patterning of the hydrogels with peptides [59,60]. This paradigm could be applied for TE-NSPs encased in micro-columns that are first crosslinked leaving unreacted functional groups that can then be bound with peptides or proteins in a secondary reaction. These molecules could influence survival, adhesion, and neurite growth, among other *in vitro* and *in vivo* outcomes. TE-NSPs could be further modified to retain or deliver growth factor and drugs, as HA hydrogels have been widely used for this purpose based on diffusion [61], protease-sensitive [62], or hydrolytic [63] degradation, on-demand force-mediated [64] and light-triggered [65–67] processes, and affinity to chemical groups [16,40,68–70]. We foresee that HA will significantly improve TE-NSPs by expanding their range of applications, design control, and prospects for successful clinical outcomes.

## Conclusions

TE-NSPs reconstruct the nigrostriatal pathway by replacing both the lost dopaminergic neurons in the nigra and their long axon fibers that innervate the striatum in their proper anatomical locations. TE-NSPs may deliver dopamine in a synaptic-based manner based on signaling from host circuitry, and may reestablish dopaminergic control over other areas of the brain needed for proper regulation of the motor circuit in the brain. In a clear contrast from common pathway reconstruction studies, TE-NSPs are fully fabricated *in vitro* for subsequent implantation without depending on *in vivo* long-distance axon growth. Moreover, TE-NSPs employ a hydrogel to encase and protect cells and axons during culture and transplantation, and in this study we sought to transition from agarose to an HA hydrogel. HA can greatly enhance our control over the mechanical and biochemical properties of the encasement, and expand its functionalities to improve *in vivo* outcomes. We confirmed that a tubular HA hydrogel could be constructed to encase a collagen/laminin core and an aggregate of rat dopaminergic neurons that extended long axon tracts and released dopamine upon stimulation. MeHA benefited neurite growth lengths and rates and elicited a reduced host inflammatory response after implantation when compared to agarose. MeHA-encased TE-NSPs could be implanted in rat brains to span the nigrostriatal pathway, maintained their integrity after implantation, and survived at least acutely. Thus, these new TE-NSPs fulfilled and at times exceeded the key structural, functional, and transplantation requirements, which supports MeHA and possibly other HA hydrogels becoming the biomaterial component of TE-NSPs moving forward. Current and future research will be centered on employing human stem cell-derived dopaminergic neurons, testing the therapeutic potential of human TE-NSPs in parkinsonian rats, scaling up the dimensions and axon lengths for use in humans, and further tuning the HA encasement to control its properties and functionalities.

## Acknowledgements

### Funding

Financial support was provided by the Department of Veterans Affairs [Merit Review I01-BX003748 (Cullen) & Career Development Award #IK2-RX002013 (Chen)], the National Institutes of Health [BRAIN Initiative U01-NS094340 (Cullen)], and the National Science Foundation [Graduate Research Fellowship DGE-1845298 (Gordián-Vélez, Galarraga)], and the Michael J. Fox Foundation [Therapeutic Pipeline Program 9998.01 (Cullen)]. Any opinion, findings, and conclusions or recommendations expressed in this material are those of the authors(s) and do not necessarily reflect the views of the Department of Veterans Affairs, National Institutes of Health, National Science Foundation, or the Michael J. Fox Foundation.

### Author contributions

W.G.V. designed the experiments, fabricated the TE-NSPs, carried out imaging, FSCV recordings, quantification, histology, and data analysis, prepared the figures, and wrote the manuscript. D.K.C. conceptualized the TE-NSP platform and directly supervised the experiments, analysis, and manuscript preparation. K.D.B. performed the transplant surgeries, assisted with histology, and provided key feedback on experimental design. J.H.G. led the synthesis of MeHA and assisted with the characterization of hydrogel properties. R.A.E. assisted with the implementation of FSCV and the interpretation and analysis of acquired data. K.D.B., J.H.G., J.E.D., R.A.E., J.A.B., and H.I.C. contributed to manuscript preparation with critical feedback, editing, and reviewing. All authors approved the final manuscript.

### Data and materials availability

All the data supporting the results and conclusions of this manuscript are available upon request.

